# Phosphorylation and lipid droplets drive the irreversible phase transition of α-Synuclein in living cells

**DOI:** 10.64898/2026.09.22.753626

**Authors:** Walid Idi, Razan Sheta, Maxime Teixeira, Abid Oueslati

**Affiliations:** CHU de Québec Research Center, Axe Neurosciences, Quebec City, QC, Canada; Department of Molecular Medicine, Faculty of Medicine, Université Laval, Quebec City, QC, Canada

**Keywords:** <u>Keywords</u>: Alpha-Synuclein, Parkinson’s Disease, phase separation, lipid droplets, proteasome, protein aggregation, optogenetics, mitochondrial dysfunction, neurodegeneration, CRY2olig

## Abstract

Aberrant phase transitions of α-synuclein (α-Syn) condensates, the pathological hallmark of Parkinson’s disease (PD) and related synucleinopathies, are difficult to dissect in living cells due to limited tools for spatiotemporal control of intracellular phase behavior. Here, we use an optogenetic-based light-inducible protein aggregation (LIPA) system to drive α-Syn liquid-liquid phase separation (LLPS) in mammalian HEK-293T cells and human iPSC-derived neurons (iNeurons). Blue-light stimulation of LIPA-α-Syn induces dynamic, liquid like biomolecular condensates that initially remain fully reversible, but gradually undergo condensate maturation into an irreversible, solid like state upon prolonged stimulation, consistent with a liquid to solid phase transition.

This transition is accompanied by a robust accumulation of phosphatase-resistant Ser129-phosphorylated α-synuclein (pS129), a hallmark biochemical feature of Lewy pathology in PD and related synucleinopathies. Mechanistically, PLK2-mediated phosphorylation at Ser129 promotes condensate stabilization and irreversibility, identifying the PLK2-pS129 axis as a key regulator of the conversion of dynamic α-Syn condensates into persistent assemblies. Importantly, transient liquid-like α-Syn condensates are efficiently cleared, primarily through the proteasomal degradation pathway. In contrast, mature solid-like condensates become refractory to degradation, resulting in proteostasis impairment and progressive intracellular α-Syn accumulation.

A major finding of this study is the identification of a strong link between α-Syn condensate maturation and lipid droplet (LD) biology. The formation of irreversible α-Syn condensates is associated with a marked increase in both LD abundance and size. Moreover, mature LIPA-α-Syn condensates exhibit extensive interactions with LDs, disrupting lipid homeostasis and increasing cellular vulnerability. These findings suggest that LDs are not merely passive bystanders but active participants in the pathological maturation of α-Syn condensates.

Collectively, our results provide mechanistic insight into the molecular and cellular processes that govern the transition of α-Syn condensates from reversible liquid assemblies to pathogenic solid-like aggregates. They identify Ser129 phosphorylation and lipid droplet interactions as critical determinants of condensate fate and establish aberrant condensate maturation as a potential source of neurotoxic α-Syn species that contribute to the pathogenesis of PD and other synucleinopathies.

## INTRODUCTION

Proteins can self-assemble into a spectrum of higher-order structures, ranging from highly dynamic, reversible condensates to stable and irreversible fibrils ^1–3^. These assemblies are central to normal cellular regulation but also underlie many proteinopathies, including neurodegenerative diseases ^4^.

In synucleinopathies, including Parkinson’s disease (PD), aberrant liquid-liquid phase separation (LLPS) of α-Syn is proposed to represent an early step in a pathogenic cascade that culminates in the formation of proteinaceous inclusions, termed Lewy bodies (LBs), in distinct brain regions ^3^. However, how α-synuclein (α-Syn) undergoes phase transitions within living cells, and how these transitions relate to the emergence of irreversible aggregates, remains poorly understood.

α-Syn is a 140-amino acid intrinsically disordered protein composed of an amphipathic N-terminal membrane-binding region, a hydrophobic region that promotes self-assembly, and an acidic C-terminal region that regulates molecular interactions and aggregation^5^. Under physiological conditions, α-Syn exists primarily as a natively unfolded monomer and possibly as a helical tetramer ^6–9^. In disease, α-Syn misfolds and assembles into higher-order species, progressing from soluble oligomers to insoluble amyloid fibrils that form the core of LBs ^10–12^. However, oligomeric and fibrillar species may coexist, and their formation may not follow a strictly linear sequence ^3,11,12^. This conversion of a functional protein into a toxic aggregate is a key event in PD pathogenesis, yet the molecular mechanisms and temporal sequence of these conformational transitions remain elusive.

Furthermore, LBs are not homogeneous deposits composed solely of α-Syn fibrils. In addition to α-Syn, they contain vesicles, fragmented organelles, membranes, and lipid material ^13–15^ . Cellular models further indicate that LB-like inclusions mature through the progressive recruitment and reorganization of proteins and organelles and that this maturation process may contribute to cellular dysfunction ^14,16,17^ . These findings support the view that LBs develop through multiple structural and biochemical states rather than through the simple accumulation of terminal amyloid fibrils.

Recent advances in the biology of biomolecular condensates and LLPS have transformed our understanding of protein aggregation in neurodegeneration ^18–24^. LLPS provides a fundamental mechanism by which cells organize biochemical reactions into membraneless compartments ^22,23^. Mounting evidence suggests that pathological protein aggregation, including α-Syn aggregation, can proceed via transient, liquid-like condensate intermediates that gradually lose their dynamic properties and transform into irreversible amyloid structures ^25–28^. This emerging view implies that protein aggregates are not static end products but dynamic assemblies that mature through discrete biophysical states ^18,27^. This emerging view implies that protein aggregates are not static but dynamic assemblies that mature through discrete biophysical states ^27–29^. However, how this physical maturation is coupled to biochemical remodeling in living cells remains unclear.

One of the most prominent biochemical hallmark of Lewy pathology is phosphorylation of α-Syn at Ser129 (S129) ^30,31^ . Its mechanistic role remains controversial because pS129 has been reported to promote, inhibit, or follow α-Syn aggregation depending on the experimental model. In seeded model, pS129 accumulates after initial protein aggregation and can inhibit further fibril formation and toxicity, suggesting that it may mark aggregate maturation rather than initiate it ^32^. Polo-like kinase 2 (PLK2) phosphorylates α-Syn at Ser129 but can also influence α-Syn turnover through autophagic pathways ^33–35^ . Thus, the temporal relationship between pS129 accumulation and condensate persistence must be established.

Lipids provide another potential interface between α-Syn condensation and aggregate maturation. The α-Syn N-terminus part binds lipid membrane, and membrane composition can either promote or inhibit its self-assembly ^36–38^ . Lipid droplets (LDs) are neutral-lipid storage organelles that regulate lipid availability and cellular adaptation to metabolic stress ^39–41^. α-Syn can associate with LD surfaces, and abnormal LD accumulation can promote formation of proteolysis-resistant α-Syn species ^42,43^. Recent work further showed that increasing abundance of LD promotes the formation of LD-rich α-Syn condensates that sequester LDs ^44^. However, it remains unclear how LD accumulation evolves during controlled condensate maturation and whether LD availability influences the transition from reversible to irreversible α-Syn aggregates.

A major barrier to dissecting these transitions in living cells has been the lack of tools that afford precise control over intracellular phase behavior ^45–47^. Preformed-fibril paradigms reproduce seeding and inclusion formation but bypass initial nucleation, whereas overexpression models generally produce aggregates slowly and asynchronously ^48,49^. To overcome the limitations of conventional aggregation models, we employed the Light-Inducible Protein Aggregation (LIPA) system ^50,51^. This optogenetic approach uses a fusion of Arabidopsis cryptochrome-2 oligomerization domain (CRY2olig), which undergoes rapid and reversible clustering upon blue-light illumination, with mCherry and human α-Syn ^52^. Upon illumination, CRY2olig oligomerizes and clusters α-Syn, thereby nucleating rapid condensate formation ^50,51^. Critically, LIPA enables tight temporal control (condensates form within seconds), spatial control (single-cell and subcellular resolution), and control over stimulation duration ^50,51^, allowing us to induce α-Syn condensation and monitor its maturation with unprecedented resolution in living cells.

Using this system, we delineate the sequence of α-Syn phase transitions from reversible to irreversible condensates in mammalian cells. We identify Ser129 phosphorylation and interactions with lipid droplets (LDs) as key regulators of this process and implicate them in the formation of toxic α-Syn species relevant to synucleinopathies pathogenesis.

### Material and methods

### Cell culture and neuronal differentiation

#### Mammalian cell lines

HEK-293T cells (ATCC, Cat: CRL-3216) were cultured in Dulbecco’s Modified Eagle Medium (DMEM) (Sigma, Cat. D5706) supplemented with 10% fetal bovine serum (FBS) (Sigma, Cat. F1051) and 1% penicillin/streptomycin (ThermoFisher, Cat. 15-140-122). Cells were maintained in a humidified incubator at 37 °C under 5% COO. All cells and their engineered derivates were tested for mycoplasma every 6 months using a Mycoplasma Pro PCR Detection Kit (ABM, cat. G239).

#### Human Induced Pluripotent Stem Cells (hiPSCs) and neuronal differentiation

The hiPSC lines used in this study were derived from the parental line AIW002-02. These hiPSC lines were provided by Dr Thomas Durcan and Dr Edward Fon from the Early Drug Discovery Unit (EDDU) at The Neuro, McGill University.

The generation of these lines has been previously described and subjected to rigorous quality controls, including immunofluorescence-based pluripotency assessment, tri-lineage differentiation potential, and genomic integrity analysis using the hiPSC Genetic Analysis Kit (STEMCELL Technologies, cat. no. 07550) ^53,54^. The use of hiPSCs in this study was approved by the CHU de Québec Research Centre (#2022-6079).

hiPSCs were cultured and maintained under standard conditions as previously described ^55,56^, in mTeSR™ Plus medium (STEMCELL Technologies, cat. no. 100-0276) on Matrigel-coated plates (Corning, cat. no. 354277). Cells were passaged using an EDTA-based 0.5 M dissociation reagent.

For neuronal differentiation, the AIW002-02 hiPSC line was genetically modified to express the transcription factor NGN2 under a doxycycline-inducible system, as previously described ^55,56^. Briefly, cells were transduced with lentiviral vectors encoding NGN2 and rtTA, allowing for rapid and controlled neuronal conversion upon exposure to doxycycline. Following transduction, NGN2-inducible hiPSCs were expanded and maintained under standard culture conditions until use.

Neuronal induction began one day after initial seeding, defined as day in vitro 0 (DIV 0), by replacing the culture medium with Day 0/1 medium consisting of DMEM/F12 supplemented with N2, B27, non-essential amino acids (NEAA), BDNF, GDNF, and mouse laminin, containing 2 µg/mL doxycycline to induce NGN2 expression. At DIV 1, the medium was completely renewed.

At DIV 2, cells were enzymatically dissociated with Accutase and re-seeded onto glass coverslips pre-treated with poly-L-ornithine (50 µg/mL) and laminin (10 µg/mL). The medium was replaced with Day 2 medium, based on Neurobasal supplemented with N2, B27, GlutaMAX, NEAA, BDNF, GDNF, mouse laminin, and 2 µg/mL doxycycline. Neurons were then maintained by half-medium changes every two days, with continuous doxycycline induction, until experimental use (typically between DIV 14 and DIV 18).

### Plasmid Cloning

The pcDNA plasmid encoding α-Syn-Cry2olig-mCherry (LIPA-α-Syn) was generated and described in our previous work ^57^. The point mutation S129A in α-Syn was introduced into the α-Syn-Cry2olig-mCherry backbone by whole-plasmid amplification using overlapping mutagenic primers, enabling site-specific incorporation of the mutation.

The plasmids YFP-CL1 (Addgene, cat. no. 11950), a fusion protein containing a short degron targeting the proteasome, and Ub-R-YFP (Addgene, cat. no. 11948), an unstable substrate degraded according to the N-end rule ^58^, were used as reporters of ubiquitin-proteasome system (UPS) activity.

The human PLK2 plasmid was kindly provided by Dr. Hilal Lashuel (Brain and Institute, EPFL, Switzerland).

The PiggyBac vector containing LIPA-α-SYN was obtained from the vector PiggyBac-TA-ERN (Addgene, cat. 80474). The cloning strategies were achieved via standard molecular cloning methods, Gateway® BP and LR Clonase™ reactions, following the manufacturer’s instructions (Thermo Fisher Scientific, cat. 11789020 and 11791020).

The integrity of all plasmids was verified by PCR and by sequencing of the whole insert for all generated constructs.

### Stable Expression of LIPA-α-Synuclein construct

#### Construction of the LIPA-α-synuclein lentiviral expression vector

The LIPA-α-Syn construct was generated using the lentiviral vector pLenti CMV Puro DEST (w118-1) (Addgene, cat. no. 17452). The LIPA-α-Syn coding sequence was cloned into the pLenti backbone by restriction enzyme-based cloning.

We used a PiggyBac transposon-based delivery system to stably express the LIPA module in our NGN2-hiPSC line. iPSCs were transfected with the PiggyBac vector(s) (WT LIPA-α-SYN), and a hyperactive PiggyBac transposase plasmid (pCMV-hyPBase) (a gift from the Canadian Neurophotonics Platform-CERVO), using Mirus TransIT®-LT1 Transfection Reagent (Mirus, cat. MIR 2304) as previously described ^59^. Following antibiotic selection G418 at 200 µg/ml, integration was confirmed via protein expression levels. Plasmid DNA was isolated and verified by restriction digest analysis and Sanger sequencing to confirm correct insertion, orientation, and integrity of the LIPA-α-syn cassette.

#### Production and packaging of lentiviral particles

Lentiviral particles were produced by transient transfection of HEK-293T cells. Cells were co-transfected with the transfer plasmid pLenti LIPA-α-Syn, the packaging plasmid pCMV-R8.74, and the envelope plasmid pMD2.G (VSV-G) using a standard calcium phosphate transfection protocol, with plasmids transfected at an optimized ratio to ensure efficient viral production. After transfection, cells were maintained in complete culture medium, and viral supernatants were collected at 48h and 72h post-transfection. Supernatants were clarified by low-speed centrifugation and filtered through 0.45 µm filters to remove cellular debris, and viral preparations were used immediately for downstream applications. Viral titers were estimated through qPCR using a lentiviral titration kit (ABM, cat. no. LV900) prior to subsequent experiments.

#### Ubiquitin-Proteasome System (UPS) Reporters

UPS activity was assessed using the fluorescent reporters YFP-CL1 and Ub-R-YFP described earlier. Plasmids encoding these reporters were transiently transfected into LIPA-α-Syn-expressing HEK-293T cells using the calcium phosphate method, as previously described ^50,51^, to assess the impact of controlled α-Syn aggregation on UPS-mediated protein degradation. Fluorescence levels were used as an inverse readout of proteasomal activity, with accumulation of signal reflecting impaired substrate degradation. As a positive control for proteasome inhibition, cells were treated with epoxomicin (100 nM, 12h), inducing robust and reproducible accumulation of reporter fluorescence.

### Induction of LIPA-α-Syn aggregation

Aggregation of the chimeric LIPA-α-Syn protein was induced by photoactivation using blue light (λ = 456 nm). Light intensity was adjusted according to cell type: 0.8 mW/mm² for HEK-293T cells and 0.1 mW/mm² for iNeurons. Cells were fixed in 4% paraformaldehyde (PFA) containing 3% sucrose for 15 min at room temperature, followed by three washes in Dulbecco’s phosphate-buffered saline (dPBS). When mentioned, to selectively remove the soluble cytosolic fraction of LIPA-α-Syn while preserving Triton-insoluble aggregated species, Triton X-100 was added directly to the fixation solution at a final concentration of 1% for HEK-293T cells and 0.5% for iNeurons to preserve neurite integrity ^60^.

### Pharmacological and Biochemical Treatments

#### Assay of liquid-liquid phase separation (LLPS) properties

To probe liquid-liquid phase separation (LLPS) properties, cells were treated with 0.625-5% 1,6-hexanediol for 10min, then fixed in 4% PFA containing 3% sucrose and 1% Triton X-100.

#### Modulation of lipid metabolism

To stimulate lipid droplet formation, cells were incubated with oleic acid (100-600 µM, complexed to BSA) for 24 or 72h.

Triglyceride synthesis was blocked by pretreatment (24h) with DGAT1 inhibitor T863 (1-10 µM) and DGAT2 inhibitor PF-06424439 (1-10 µM).

#### Inhibition of degradation pathways

The following inhibitors were added at the onset of illumination and maintained until the end of the experiment at the indicated final concentrations. To block the proteasome, cells were treated with MG132 (5 µM) and epoxomicin (EPX, 50 nM). To block autophagy, cells were treated with 3-methyladenine (3-MA, 5 mM) to block Autophagy initiation and chloroquine (CQ, 50 µM) and ammonium chloride (NHOCl, 10 mM) to block Autophagosome-lysosome fusion.

#### Alkaline Phosphatase Calf Intestinal Treatment (CIP)

Fixed HEK-293T cells (4% PFA / 3% sucrose / 1% Triton X-100) were incubated overnight at 37 °C with QuickCIP enzyme (20 units/mL) in a specific reaction buffer (50 mM Tris-HCl, 100 mM NaCl, 10 mM MgClO, pH 7.9). Following enzymatic treatment, standard immunocytochemistry was performed to quantify the residual pS129 signal.

#### Inhibition of Polo-like kinase (PLK) activity

To inhibit PLK activity, cells were pretreated with BI2536 (1 or 5 µM; cat. no. S1109, Selleck Chemicals, Houston, TX, USA) for 2 h before illumination. BI2536 was maintained during the 6 h illumination period.

### Immunocytochemistry

After fixation, cells were permeabilized with 0.25% Triton X-100 in PBS for 15 min. Blocking was then performed for 1h in a solution containing 5% normal goat serum (NGS), 1% bovine serum albumin (BSA), and 0.1% Triton X-100 in PBS. Coverslips were incubated with primary antibodies diluted in blocking buffer either for 2h at room temperature (RT) or overnight at 4 °C. After three PBS washes, cells were incubated for 1h at RT with secondary antibodies conjugated to appropriate Alexa Fluor dyes (Table 1). For HEK-293T cells, nuclei were counterstained with DAPI (4′,6-diamidino-2-phenylindole, 1:10 000). After a final PBS wash, coverslips were mounted onto slides using Fluoromount-G™ mounting medium (Electron Microscopy Sciences). Antibody details and dilutions used in this study are provided in Table 1.

**Table 1:**
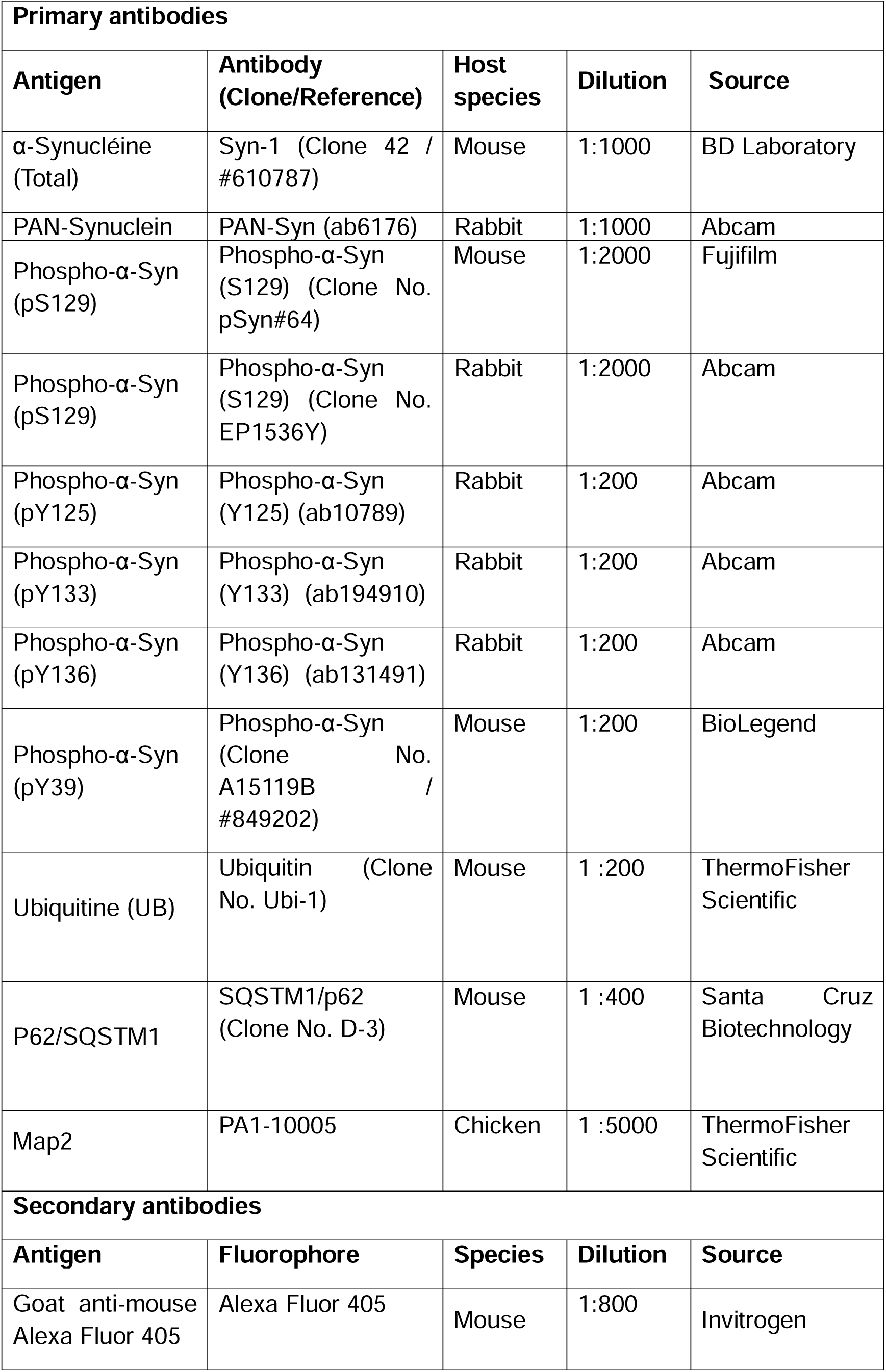
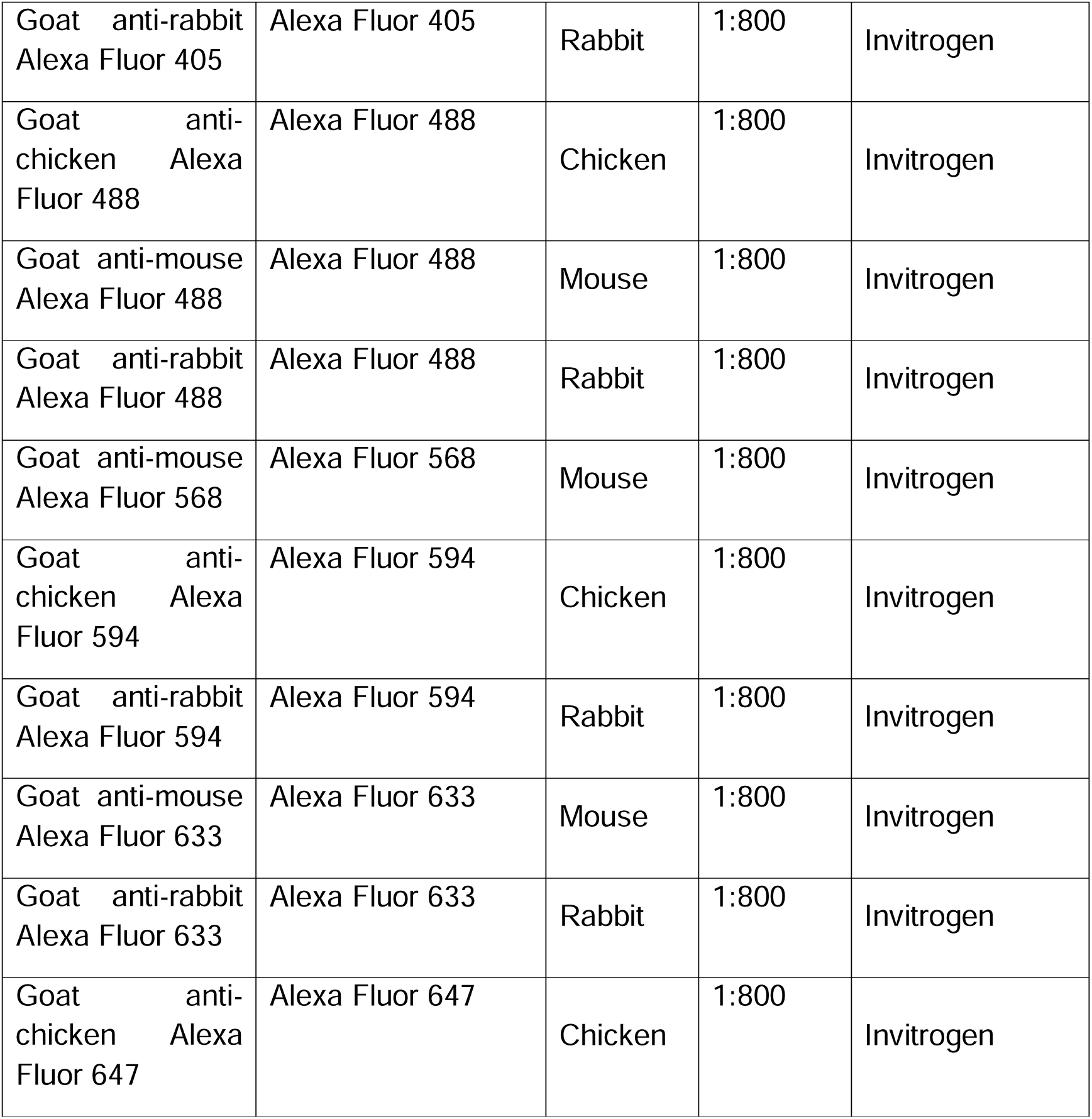
Table summarizing the antibodies used in this study.

| Primary antibodies |  |  |  |  |
| --- | --- | --- | --- | --- |
| Antigen | Antibody<br>(Clone/Reference) | Host<br>species | Dilution | Source |
| α-Synucléine<br>(Total) | Syn-1 (Clone 42 /<br>#610787) | Mouse | 1:1000 | BD Laboratory |
| PAN-Synuclein | PAN-Syn (ab6176) | Rabbit | 1:1000 | Abcam |
| Phospho-α-Syn<br>(pS129) | Phospho-α-Syn<br>(S129) (Clone No.<br>pSyn#64) | Mouse | 1:2000 | Fujifilm |
| Phospho-α-Syn<br>(pS129) | Phospho-α-Syn<br>(S129) (Clone No.<br>EP1536Y) | Rabbit | 1:2000 | Abcam |
| Phospho-α-Syn<br>(pY125) | Phospho-α-Syn<br>(Y125) (ab10789) | Rabbit | 1:200 | Abcam |
| Phospho-α-Syn<br>(pY133) | Phospho-α-Syn<br>(Y133) (ab194910) | Rabbit | 1:200 | Abcam |
| Phospho-α-Syn<br>(pY136) | Phospho-α-Syn<br>(Y136) (ab131491) | Rabbit | 1:200 | Abcam |
| Phospho-α-Syn<br>(pY39) | Phospho-α-Syn<br>(Clone No.<br>A15119B /<br>#849202) | Mouse | 1:200 | BioLegend |
| Ubiquitine (UB) | Ubiquitin (Clone<br>No. Ubi-1) | Mouse | 1 :200 | ThermoFisher<br>Scientific |
| P62/SQSTM1 | SQSTM1/p62<br>(Clone No. D-3) | Mouse | 1 :400 | Santa Cruz<br>Biotechnology |
| Map2 | PA1-10005 | Chicken | 1 :5000 | ThermoFisher<br>Scientific |
| Secondary antibodies |  |  |  |  |
| Antigen | Fluorophore | Species | Dilution | Source |
| Goat anti-mouse<br>Alexa Fluor 405 | Alexa Fluor 405 | Mouse | 1:800 | Invitrogen |
| Goat anti-rabbit<br>Alexa Fluor 405 | Alexa Fluor 405 | Rabbit | 1:800 | Invitrogen |
| Goat anti-chicken<br>Alexa Fluor 488 | Alexa Fluor 488 | Chicken | 1:800 | Invitrogen |
| Goat anti-mouse<br>Alexa Fluor 488 | Alexa Fluor 488 | Mouse | 1:800 | Invitrogen |
| Goat anti-rabbit<br>Alexa Fluor 488 | Alexa Fluor 488 | Rabbit | 1:800 | Invitrogen |
| Goat anti-mouse<br>Alexa Fluor 568 | Alexa Fluor 568 | Mouse | 1:800 | Invitrogen |
| Goat anti-chicken<br>Alexa Fluor 594 | Alexa Fluor 594 | Chicken | 1:800 | Invitrogen |
| Goat anti-rabbit<br>Alexa Fluor 594 | Alexa Fluor 594 | Rabbit | 1:800 | Invitrogen |
| Goat anti-mouse<br>Alexa Fluor 633 | Alexa Fluor 633 | Mouse | 1:800 | Invitrogen |
| Goat anti-rabbit<br>Alexa Fluor 633 | Alexa Fluor 633 | Rabbit | 1:800 | Invitrogen |
| Goat anti-chicken<br>Alexa Fluor 647 | Alexa Fluor 647 | Chicken | 1:800 | Invitrogen |

### Cell viability assay

Cells viability was assessed using 3-(4,5-dimethylthiazol-2-yl)2,5-diphenyltetrazolium bromide (MTT) assay ^61^. MTT solution (5mg/mL in PBS) was added at 25 µL per well containing 500 µL of culture medium, followed by incubation for 2 h at 37°C in the dark. The medium was then removed, and the formazan crystals were dissolved in 500 µL of acidified isopropanol (1 N HCl, 1:25, v/v). After thorough mixing, 200 µL from each well was transferred to a 96-well plate, and absorbance was measured at 570 nm using a microplate reader.

### Microscopy

#### Standard confocal imaging

Confocal microscopy images were acquired using a Zeiss LSM800 microscope (Zeiss, Oberkochen, Germany). Subsequent image processing and analysis were performed with ZEN (Zeiss) and ImageJ/Fiji software.

#### Fluorescence Recovery After Photobleaching (FRAP)

FRAP experiments were performed on a Leica TCS SP8 microscope at 37 °C in a COO-controlled chamber. A 3×3 µm region of interest (ROI) at the center or periphery of an aggregate was photobleached using a 561 nm laser at 100% intensity. Fluorescence recovery was recorded via time-lapse imaging, and recovery curves were calculated after data normalization and passive photobleaching correction.

#### STED (STimulated Emission Depletion) microscopy

Super-resolution imaging was conducted on a Leica TCS SP8 STED 3X microscope equipped with an oil-immersion objective (HC PL APO CS2 100×/1.40). Fluorescence depletion was performed with a 592 nm laser (for Alexa Fluor 488) or a 660 nm laser (for mCherry/Alexa Fluor 568). Images were acquired using a time-gating window of 0.3-3.5 ns to minimize autofluorescence and enhance resolution. Raw images were deconvolved using Huygens Professional software (Scientific Volume Imaging) to optimize nanoscale resolution, and Z-stacks were reconstructed in 3D with IMARIS software (Bitplane).

### Quantification and Statistical Analysis

#### Image Processing

Initial processing of raw (.czi) files was carried out in ZEN (Zeiss). Most quantifications (cell counts with aggregates, fluorescence intensity) were performed manually or semi-automatically in ImageJ/Fiji software. The StarDist plugin was used for nuclear segmentation and cell counting. The 3D Objects Counter plugin was used to quantify aggregate number and size, as well as aggregate-associated fluorescence intensity.

#### Automated Quantification

Lipid droplet analysis was automated using CellProfiler (Broad Institute). A segmentation pipeline adapted from Adomshick et al., (2020) was established to identify nuclei (DAPI), lipid droplets and cell boundaries, enabling quantification of lipid droplet number and size per cell. ^62^

#### Statistical Analysis

Statistical analyses were performed using GraphPad Prism v9. Data are presented as mean ± standard deviation (SD) or standard error of the mean (SEM). Statistical comparisons were made using analysis of variance (ANOVA) tests followed by appropriate post hoc tests. The significance threshold was set at p < 0.05.

## Results

### Prolonged light-induced clustering promotes the transition of α-Syn assemblies from reversible liquid-like to irreversible gel-like states

Our team has developed a novel optogenetic approach to induce α-Syn aggregation, termed the Light-Inducible Protein Aggregation (LIPA) system ^50,51^. This platform provides, for the first time, precise temporal control over the initiation of α-Syn aggregation through blue-light stimulation and promotes the formation of inclusion bodies that recapitulate key features of authentic LBs ^50^. Using this system, we observed that increasing the duration of blue-light exposure leads to the formation of aggregates with distinct stability profiles, with more stable and irreversible clusters forming as stimulation time increases. This observation suggests that the temporal resolution provided by the LIPA system can be leveraged to capture early dynamic phases of α-Syn aggregation in living cells.

To test this hypothesis, we exposed HEK-293T cells stably expressing LIPA-α-Syn to blue light (0.8 mW/mm²) for varying durations (3, 6, 12, or 24h). Aggregate stability was subsequently assessed by quantifying the proportion of cells retaining α-Syn inclusions at 0, 3, 6, 12, and 24h following cessation of illumination (post-illumination) (**Figure 1A**). We reasoned that reversible aggregates would rapidly dissipate upon light withdrawal, whereas a greater proportion of cells would retain aggregates once these structures became irreversible and persisted for several hours or longer.

**Figure 1:**
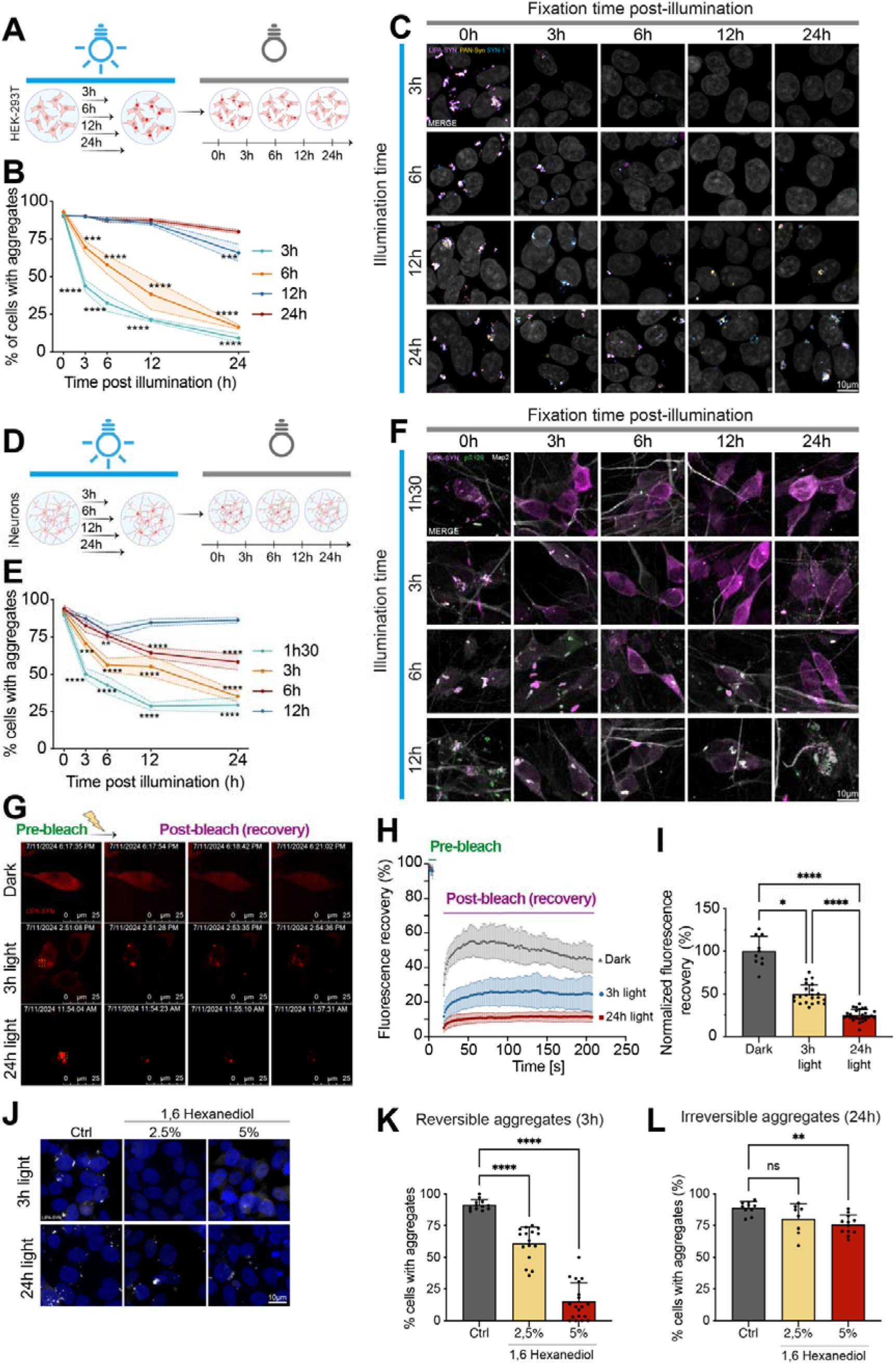
Prolonged illumination induces a phase transition of LIPA-α-Syn aggregates from a reversible to an irreversible state. (**A**) Overview of the experimental protocol used to study the stability of LIPA-α-Syn aggregates in HEK-293T cells. (**B**) Quantification of aggregate-positive cells during the post-illumination period after 3, 6, 12, and 24h of illumination. (* = vs 0h) (N = 3). Data are presented as means ± SEM; \**p* ≤ 0.05; \*\**p* ≤ 0.01; \*\*\**p* ≤ 0.001; \*\*\*\**p* < 0.0001. (**C**) Confocal images of HEK-293T cells expressing LIPA-α-Syn (magenta) exposed to blue light for 3, 6, 12, or 24h (0.8 mW/mm²) and fixed with 4% PFA - 1% Triton X-100 at various post-illumination time points (0, 3, 6, 12, and 24h). Cells were immunostained with Syn-1 (cyan) and PAN-Syn (yellow), two antibodies recognizing distinct α-Syn epitopes to exclude loss of signal due to protein truncation. Nuclei were counterstained with DAPI (gray). (Scale bar = 10 µm). (**D**) Overview of the experimental protocol used to study the stability of LIPA-α-Syn aggregates in iNeurons. (**E**) Quantification of aggregate-positive cells during the post-illumination period after 1 h 30, 3, 6, or 12h of illumination. (* = vs 0h) (N = 3). Data are presented as means ± SEM; \**p* ≤ 0.05; \*\**p* ≤ 0.01; \*\*\**p* ≤ 0.001; \*\*\*\**p* < 0.0001. (**F**) Confocal images of iNeurons expressing LIPA-α-Syn (magenta) exposed to blue light for 1h30, 3, 6, or 12h (0.1 mW/mm²) and fixed with 4% PFA - 0.5% Triton X-100 at different post-illumination time points (0, 3, 6, 12, and 24h). Neurons were immunolabeled with pSer129 (green) and MAP2 (gray) antibodies. (Scale bar = 10 µm). (**G**) Representative FRAP images from live-cell imaging experiments showing HEK-293T cells expressing LIPA-α-Syn (red), either non-exposed (0h) or exposed to blue light for 3h or 24h (0.8 mW/mm²). Images were acquired (1) before photobleaching (pre-bleach), (2) 20s, (3) ∼2min, and (4) ∼3min after photobleaching (post-bleach/recovery). (Scale bar = 25 µm). (**H**) Quantification of fluorescence recovery over time (non-normalized) (N = 3). (**I**) Quantification of fluorescence recovery at the last acquisition time point (normalized) (N = 3). Data are presented as means ± SD; \**p* ≤ 0.05; \*\**p* ≤ 0.01; \*\*\**p* ≤ 0.001; \*\*\*\**p* < 0.0001. (**J**) Confocal images of HEK-293T cells expressing LIPA-α-Syn (gray) exposed to blue light for 3 or 24h (0.8 mW/mm²), treated DMSO or with 2.5 or 5% 1,6-hexanediol, and fixed with 4% PFA - 1% Triton X-100. Nuclei were stained with DAPI (gray). (Scale bar = 10 µm). (**K**) Quantification of aggregate-positive cells LIPA-α-Syn (gray) after 3h of illumination ± 1,6-hexanediol, after fixation with 4% PFA / 1% Triton X-100. (* = vs DMSO) (N = 3). Data are presented as means ± SD; \**p* ≤ 0.05; \*\**p* ≤ 0.01; \*\*\**p* ≤ 0.001; \*\*\*\**p* < 0.0001. (**L**) Quantification of aggregate-positive cells after 24h of illumination ± 1,6-hexanediol, after fixation with 4% PFA / 1% Triton X-100. (* = vs DMSO) (N = 3). Data are presented as means ± SD; \**p* ≤ 0.05; \*\**p* ≤ 0.01; \*\*\**p* ≤ 0.001; \*\*\*\**p* < 0.0001.

Post-illumination analysis demonstrated that short exposure times (3h and 6h) predominantly generated reversible clusters, as the number of cells harboring aggregates dramatically decreased after the light was turned off, reaching near-complete clearance 24h post-illumination (9% and 16% of remaining cells with aggregates, for 3 and 6h of light exposure, respectively) (**Figure 1B, C**). Consistent with this rapid clearance, the proportion of cells harboring aggregates was significantly reduced at post-illumination time points compared with 0h (**Figure 1B**). In the other hand, prolonged exposure to blue light for 12h and 24h resulted in the formation of markedly more stable inclusions, with more than 70% - 80% of cells retained aggregates 24h post-illumination (**Figure 1B, C**). Together, these results confirmed that increasing the duration of the blue light stimulation correlates with a transition from reversible to irreversible α-Syn cluster states in living cells.

Importantly, similar results were observed in iPSC-derived human neurons (iNeurons), in which a shorter illumination range (1.5-12h) was used to account for the distinct aggregation kinetics in this cellular model and capture the transition from reversible to persistent α-Syn assemblies. Inclusions generated by a short exposure to blue light (1.5h and 3h) were predominantly reversible aggregates, with a rapid and significant decrease in the percentage of positive neurons within 24h after the end of illumination compared with 0h (30% and 35%) (**Figure 1D-F**). At 6h of illumination, aggregates entered a transition phase: the proportion of positive neurons remained high and relatively stable between 0h and 3h (∼94% to ∼82%), then declined more gradually to ∼58% at 24h. After 12h of illumination, the proportion of positive cells remained almost unchanged, around 85-92% between 0h and 24h (**Figure 1D-F**).

Next, we investigated the molecular dynamics of the reversible and irreversible α-Syn aggregates using fluorescence recovery after photobleaching (FRAP). This approach makes it possible to estimate the mobile fraction and the rate of turnover of molecules within an assembly, parameters classically used to distinguish “liquid-like” states from more rigid structures ^22^. Reversible aggregates formed after 3h of illumination retain a certain molecular dynamic, although distinct from that of the diffuse protein. Indeed, we observe a significant decrease in recovery compared with the Dark condition, consistent with a condensed assembly exhibiting increased viscosity without complete diffusion blockade ^22,28^ (**Figure 1G-I**). In contrast, after 24h of illumination, recovery decreases significantly compared with Dark and 3h conditions, indicating a major loss of mobility and a progressive rigidification of the inclusions (**Figure 1G-I**). These results support a transition from a more dynamic state (3h) to a solid state (24h), often described as the maturation of α-Syn assemblies ^3^. This difference in dynamics is consistent with the differential sensitivity to 1,6-hexanediol, a compound that disrupts the weak hydrophobic interactions typical of LLPS ^63^. Early assemblies (3h) show a dose-dependent dissociation with a significant effect at 2.5% and becomes almost complete at 5% 1,6-hexanediol, confirming that assemblies are maintained by weak and labile interactions (**Figure 1J & K**). In contrast, mature assemblies (24h) are largely resistant to 1,6-hexanediol. Even at 5%, approximately 75% of the cells are retaining aggregates, confirming that temporal maturation locks inclusions into a solid state that is mostly insensitive to perturbation of weak interactions (**Figure 1L**).

### Phosphatase-resistant Ser129 phosphorylation marks α-Syn irreversible aggregates, while PLK2-induced phosphorylation enhances aggregates stability

Given the strong association between S129 phosphorylation (pS129) and the accumulation of pathological α-Syn aggregates ^64^, we sought to determine how this post-translational modification (PTM), as well as phosphorylation at other disease-relevant residues (Y39, Y125, Y133, and Y136), evolves during α-Syn phase transitions in living cells. To address this question, we characterized the temporal dynamics of pS129, pY39, pY125, pY133, and pY136 as α-Syn progressed from a soluble state to condensed assemblies and ultimately to aggregated species in both HEK-293T cells and iNeurons.

We observed a progressive increase in α-Syn phosphorylation at most residues as a function of illumination time. Among the sites examined, pS129 displayed the most rapid and robust response in both HEK-293T cells (**Figure 2 A, B**) and iNeurons (**Figure 2 C, D**). The proportion of cells harboring pS129-positive aggregates increased from virtually undetectable levels to approximately 86% in HEK-293T cells and 70% in iNeurons within just 3h of light stimulation, with the increase reaching statistical significance at subsequent illumination time points and ultimately reaching ∼97-100% at later time points. Other disease-associated phosphorylation sites, particularly Y39 and Y125, also exhibited significant illumination-dependent increases, although to a lesser extent than pS129 in both cellular models (**Figure 2A–D; Figure S1**).

**Figure 2:**
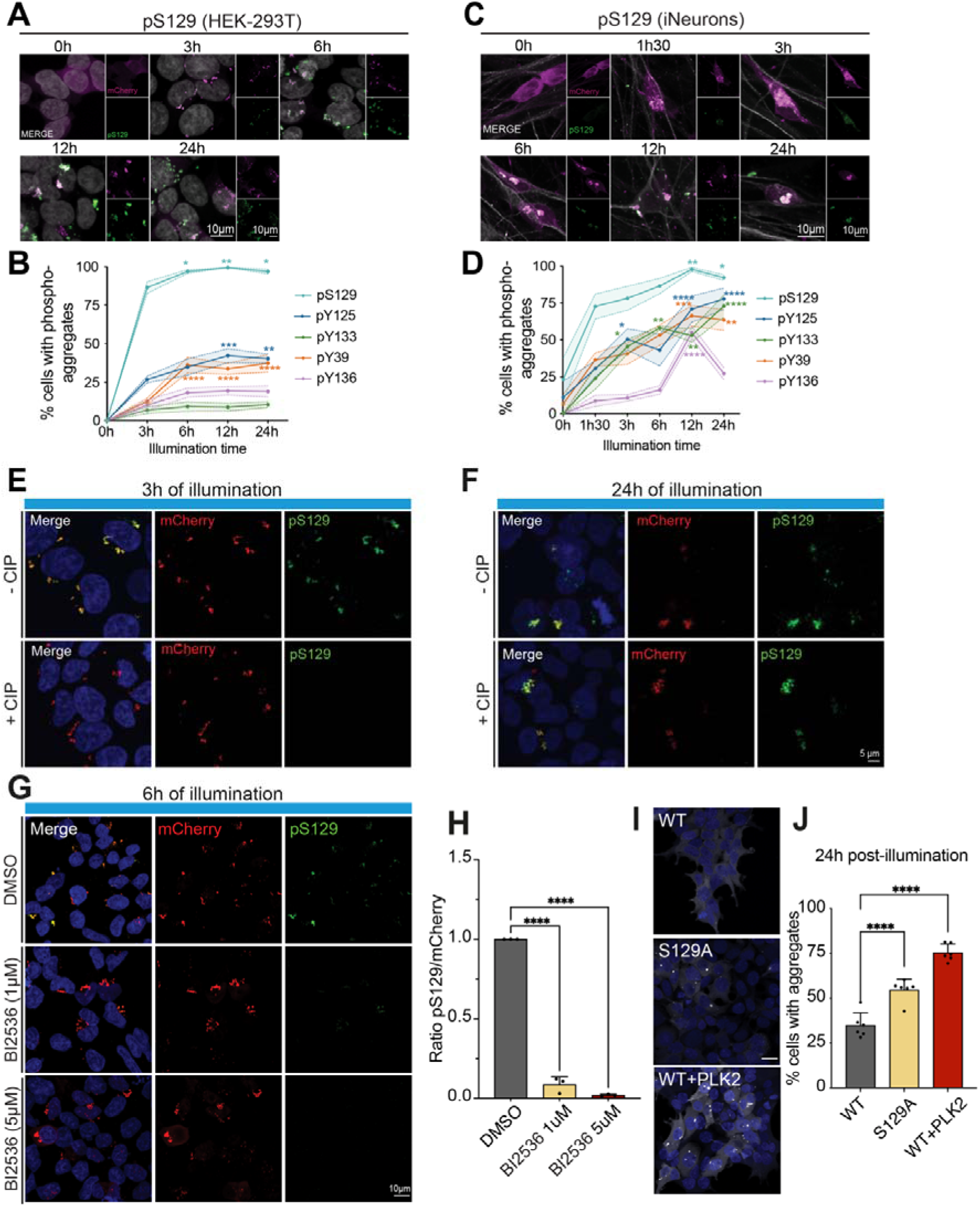
Phosphorylation at Ser129 is important for the transition of LIPA-α-Syn aggregates from a reversible to an irreversible state. (**A**) Confocal images of HEK-293T cells expressing LIPA-α-Syn (magenta) exposed to blue light for 3, 6, 12, or 24h (0.8 mW/mm²) or not exposed (0h) and fixed with 4% PFA. Cells were immunolabeled for pS129 PTM (green) (Scale bar = 10 µm). (**B**) Quantification of the percentage of LIPA-α-Syn aggregate-positive cells for each PTM, normalized to the total number of cells containing aggregates. Values were compared with the 3 h condition. (* = vs 3h) (N = 3). Data are presented as means ± SEM; \**p* ≤ 0.05; \*\**p* ≤ 0.01; \*\*\**p* ≤ 0.001; \*\*\*\**p* < 0.0001. (**C**) Confocal images of iNeurons expressing LIPA-α-Syn (magenta) exposed to blue light for 1 h 30, 3, 6, 12, or 24 h (0.1 mW/mm²) or not exposed (0 h) and fixed with 4% PFA. Cells were immunolabeled for pS129 PTM (green) (Scale bar = 10 µm). (**D**) Quantification of the percentage of LIPA-α-Syn aggregate-positive cells for each Lewy body-associated marker relative to the total number of cells containing aggregates. Values were compared with those for the 3 h condition, which represents the most reversible aggregates. (* = vs 3 h) (N = 3). Data are presented as means ± SEM; *p ≤ 0.05; **p ≤ 0.01; ***p ≤ 0.001; ****p < 0.0001. (**E**) & (**F**) Confocal images of HEK-293T cells expressing LIPA-α-Syn (red) exposed to blue light for 3h or 24h (0.8 mW/mm²), fixed with 4% PFA - 1% Triton X-100, with or without CIP treatment. Cells were immunolabeled with pS129 (Abcam, green) to assess resistance or sensitivity to CIP-mediated dephosphorylation and stained with DAPI (blue). (**G**) Confocal images of HEK-293T cells expressing LIPA-α-Syn (red) treated with BI2536 for 2 h (1 or 5 µM or treated with DMSO and subsequently exposed to blue light for 6h (0.8 mW/mm²) in the presence of BI2536. Cells were fixed with 4% PFA - 1% Triton X-100 and immunolabeled with pS129 (Abcam, green) to evaluate pS129 phosphorylation; nuclei were stained with DAPI (blue). (**H**) Quantification of the integrated density of the pS129 signal relative to the LIPA-α-Syn (mCherry) signal (* = vs NT) (N = 3). Data are presented as means ± SEM; \**p* ≤ 0.05; \*\**p* ≤ 0.01; \*\*\**p* ≤ 0.001; \*\*\*\**p* < 0.0001. (**I**) Confocal images of HEK-293T cells expressing the phospho-mutant variant of LIPA-α-Syn (S129A) or co-expressing LIPA-α-Syn WT with the kinase PLK2. Images were acquired after a light-induction phase followed by 24h in the dark to assess aggregate stability (gray). Nuclei were stained with DAPI (blue) (Scale bar = 10 µm). (**J**) Quantification of the percentage of cells retaining aggregates 24h after cessation of illumination (24h post-illumination) for each condition (WT, S129A, WT+PLK2) (N = 3) (* = vs WT). Data are presented as means ± SEM; \**p* ≤ 0.05; \*\**p* ≤ 0.01; \*\*\**p* ≤ 0.001; \*\*\*\**p* < 0.0001.

To distinguish pS129 associated with reversible condensates from that associated with irreversible inclusions, cells were treated with Alkaline Phosphatase Calf Intestinal (CIP). This approach was based on previous reports demonstrating that pS129 incorporated into mature α-Syn aggregates is largely resistant to phosphatase-mediated dephosphorylation ^65^. Consistent with this observation, the pS129 signal detected in reversible LIPA-α-Syn condensates formed after 3h of illumination was readily abolished by CIP treatment. In contrast, pS129 associated with irreversible aggregates generated following prolonged illumination (24h) remained largely resistant to dephosphorylation (**Figure 2E, F**).

Given the rapid and robust accumulation of pS129 during LIPA-α-Syn aggregation, we next sought to determine the kinase responsible for this phosphorylation event. Cells were treated with BI2536, a potent inhibitor of Polo-like kinases (PLKs), including PLK2, the major α-Syn Ser129 kinase ^33,35^. BI2536 treatment produced a robust, and a significant dose-dependent reduction in pS129 levels normalized to LIPA-α-Syn expression (**Figure 2G, H**), supporting a major contribution of PLK activity to S129 phosphorylation during LIPA-α-Syn phase transitions in our cellular model.

We next investigated the functional contribution of S129 phosphorylation to aggregate stabilization and the transition toward irreversible aggregation. To enhance endogenous S129 phosphorylation, PLK2 was overexpressed in cells expressing LIPA-α-Syn. In parallel, a phospho-deficient mutant, LIPA-α-Syn-S129A, was used to assess the consequences of abolishing phosphorylation at this residue. Aggregate stability was quantified by measuring the proportion of cells retaining inclusions for 24h after a 6h illumination period. PLK2 overexpression significantly increased inclusion stability, with approximately 75% of cells retaining aggregates compared with 35% in cells expressing LIPA-α-Syn alone (**Figure 2I, J**). Surprisingly, the phospho-deficient S129A mutant also exhibited a significant increase in aggregate stability, although the effect was less pronounced than that observed following PLK2 overexpression. Together, these findings suggest that perturbation of the normal dynamic regulation of S129 phosphorylation, either by enhancing or abolishing it, promotes α-Syn aggregate stabilization.

Of note, the phase transition of LIPA-α-Syn was also accompanied by the progressive acquisition of canonical LBs markers, including ubiquitin and p62, in an illumination time-dependent manner (**Figure S2**). These findings further indicate that the transition to irreversible aggregates involves extensive biochemical remodeling and maturation rather than a simple physical liquid-to-solid transformation.

### Proteasomal clearance controls aggregate reversibility and is compromised during α-Syn phase transition

Next, we sought to determine how reversible LIPA-α-Syn inclusions dissipate following cessation of light stimulation. Specifically, we asked whether their disappearance results from passive disassembly or from active clearance by cellular protein quality-control pathways ^52^.

To address this question, HEK-293T cells stably expressing LIPA-α-Syn were treated with inhibitors of the two major protein degradation pathways. The ubiquitin-proteasome system (UPS) was inhibited using epoxomicin (EPX) or MG132, whereas the autophagy-lysosomal pathway (ALP) was blocked using ammonium chloride (NHOCl), chloroquine (CQ), or 3-methyladenine (3-MA) ^66–70^. Cells were exposed to blue light for 6h in the presence of these inhibitors to induce irreversible LIPA-α-Syn aggregation. Aggregate stability was then assessed at 0, 6, and 12h following light cessation by quantifying the proportion of cells retaining LIPA-α-Syn inclusions (**Figure 3A**).

**Figure 3:**
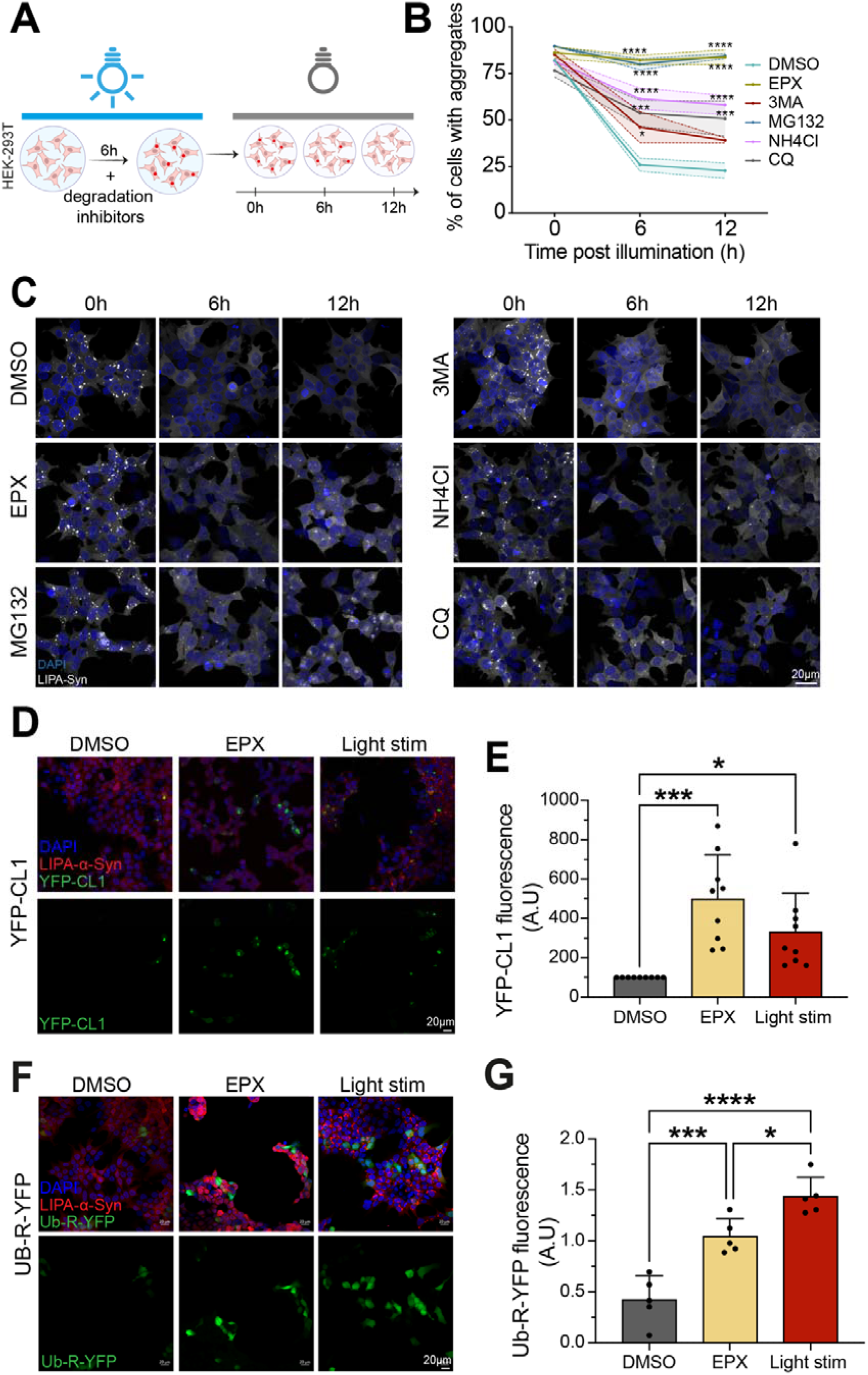
Proteasome inhibition promotes stabilization of reversible aggregates, whereas formation of irreversible aggregates compromises proteasome activity. (**A**) Overview of the experimental protocol used to study the stability of LIPA-α-Syn aggregates in HEK-293T cells following inhibition of cellular degradation pathways. (**B**) Quantification of the number of cells with LIPA-α-Syn aggregates relative to the total number of mCherry-positive cells (* = vs 0h) (N = 3). Data are presented as means ± SEM; *p ≤ 0.05; \*\**p* ≤ 0.01; \*\*\**p* ≤ 0.001; \*\*\*\**p* < 0.0001. (**C**) Confocal images of HEK-293T cells expressing LIPA-α-Syn (gray) treated with DMSO or the indicated degradation inhibitors (EPX, MG132, 3-MA, NHOCl, or CQ), exposed to blue light for 6h (0.8 mW/mm²), and fixed with 4% PFA at 24h post-illumination. (**D**) Confocal images of HEK-293T LIPA-α-Syn cells expressing YFP-CL1 (green), and LIPA-α-Syn (red) with or without exposure to blue light (12h) and with or without the degradation inhibitor EPX; cells were fixed with 4% PFA. (**E**) Quantification of YFP-CL1 fluorescence (A.U) per field (N = 3). Data are presented as means ± SEM; \**p* ≤ 0.05; \*\**p* ≤ 0.01; \*\*\**p* ≤ 0.001; \*\*\*\**p* < 0.0001. (**F**) Confocal images of HEK-293T LIPA-α-Syn cells expressing Ub-R-YFP (green) with or without exposure to blue light (12h) and with or without the degradation inhibitor EPX; cells were fixed with 4% PFA. (**G**) Quantification of Ub-R-YFP fluorescence (A.U) per field (N = 3). Data are presented as means ± SEM; \**p* ≤ 0.05; \*\**p* ≤ 0.01; \*\*\**p* ≤ 0.001; \*\*\*\**p* < 0.0001.

At the end of the illumination period (0h post-illumination), cells treated with UPS or ALP inhibitors exhibited levels of aggregation comparable to those of control cells, indicating that inhibition of protein degradation pathways does not affect aggregate formation (**Figure 3B, C**). In contrast, marked differences emerged during the recovery phase. Under control conditions, the proportion of cells containing aggregates declined rapidly at 6h and 12h after light withdrawal, consistent with the reversible nature of LIPA-α-Syn assemblies. However, inhibition of the UPS with either EPX or MG132 significantly impaired aggregate clearance, with >80% of cells still harboring inclusions 12h after light cessation (**Figure 3B, C**). Inhibition of the ALP also significantly impaired aggregate clearance, although to a lesser extent, with approximately 50-58% of cells retaining aggregates following NHOCl or CQ treatment and ∼40% following 3-MA treatment (**Figure 3B, C**). These findings demonstrate that reversible LIPA-α-Syn aggregates are actively eliminated by cellular protein quality-control mechanisms, with the UPS serving as the predominant clearance pathway. More importantly, they suggest that impairment of degradative pathways during α-Syn phase transition may contribute directly to the persistence and stabilization of irreversible inclusions.

We next investigated whether proteasomal function becomes compromised during LIPA-α-Syn phase transition. To monitor UPS activity, we employed two established proteasomal reporters, YFP-CL1 and Ub-R-YFP ^58^, in cells expressing LIPA-α-Syn. Both reporters contain degron sequences that target them for rapid proteasomal degradation; consequently, reporter accumulation and increased fluorescence indicate impaired UPS activity ^58^. Under basal conditions, prior to the induction of LIPA-α-Syn aggregation, reporter fluorescence was virtually undetectable, consistent with efficient proteasomal degradation. As expected, treatment with epoxomicin, used as a positive control, resulted in a significant and robust reporter accumulation. Strikingly, induction of light-driven α-Syn aggregation likewise caused substantial accumulation of both reporters, indicating that LIPA-α-Syn aggregate formation is sufficient to impair proteasomal activity (**Figure 3D-G**).

Collectively, these findings reveal a dynamic and self-reinforcing pathological cycle in which nascent α-Syn aggregates impair proteasomal function, thereby reducing their own clearance and promoting the stabilization of increasingly persistent assemblies. This feed-forward mechanism provides a potential explanation for how initially reversible α-Syn condensates progressively transition into irreversible proteotoxic aggregates.

### Lipid droplets accumulate during LIPA-α-Syn phase transition, recruit α-Syn aggregates and stabilize them

Analysis of the morphology of stable LIPA-α-Syn aggregates following prolonged light stimulation revealed the formation of distinctive annular structures (**Figure 4A**). Confocal imaging of cells exposed to light for 24h revealed mCherry-positive ring-like morphology with a hollow-appearing center (**Figure 4A**). Both LIPA-α-Syn and pS129 signals were predominantly enriched at the aggregate periphery, forming a shell of phosphorylated α-Syn surrounding a central region largely devoid of α-Syn signal. Notably, this morphology closely resembled previously described patterns of α-Syn accumulation surrounding LDs ^44,71,72^. To assess this possibility, cells were co-stained with LipidSpot, a lipid droplet marker, which revealed a strong spatial association between the central regions of the annular aggregates and LDs (**Figure 4B**). Line-scan intensity analysis further demonstrated a clear spatial complementarity between the signals, with peripheral LIPA-α-Syn and pS129 fluorescence flanking a central LipidSpot-positive signal (**Figure 4C, D**). STED acquisitions followed by 3D reconstructions confirm that the aggregate forms a shell enveloping the LD (**Figure S3**). Together, these findings indicate that prolonged light stimulation promotes the formation of stable α-Syn inclusions characterized by a peripheral shell enriched in phosphorylated α-Syn and surrounding LDs.

**Figure 4:**
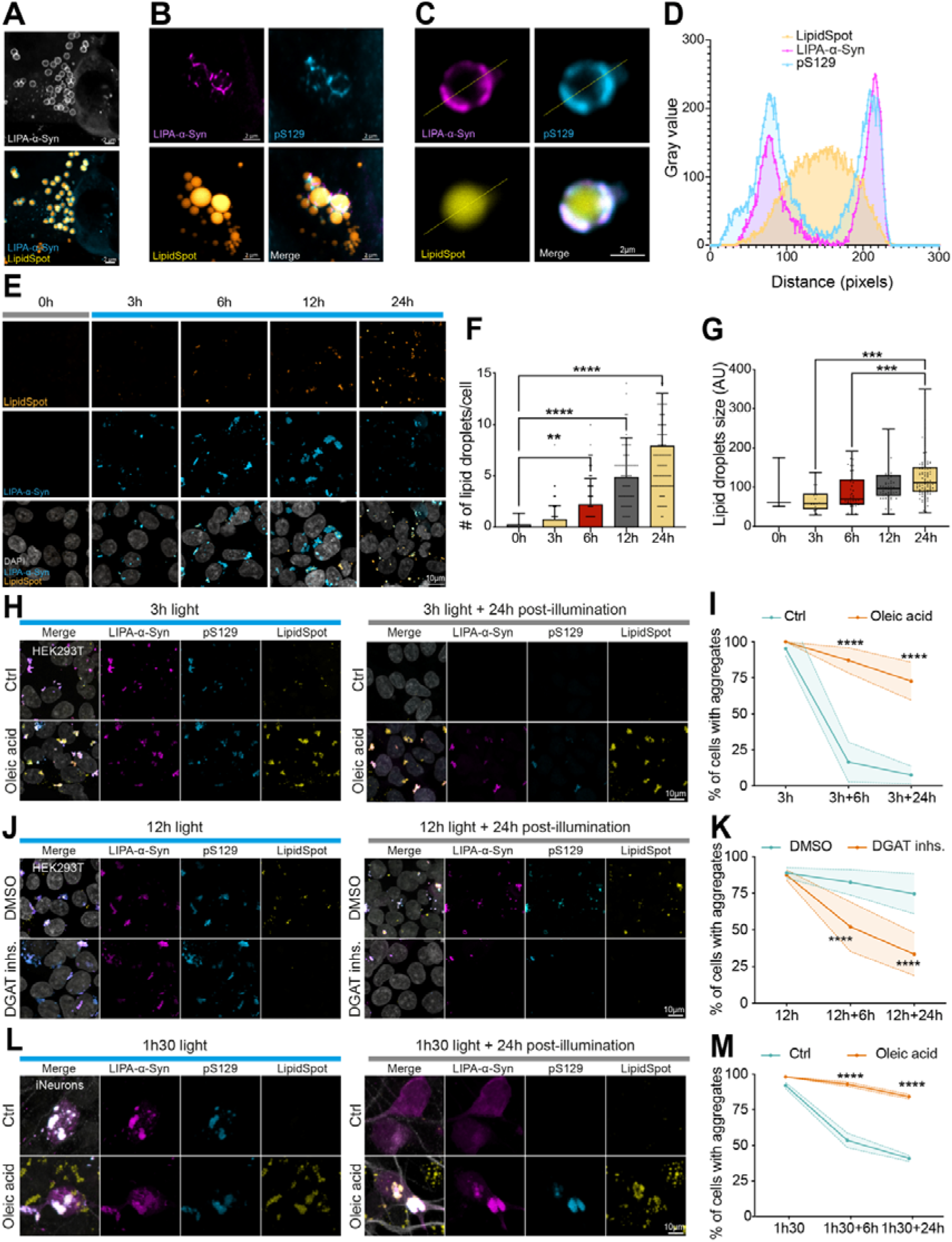
Lipid droplets play an important role in the stabilisation and persistence of α-Syn inclusions during their transition toward an irreversible state. (**A**) Confocal images of HEK-293T cells expressing LIPA-α-Syn (Top panel: gray; Bottom panel: cyan) exposed to blue light for 24 h (0.8 mW/mm²) and fixed with 4% PFA, showing accumulation of lipid droplets (LipidSpot staining, yellow). (Scale bar = 2 µm). (**B**) Confocal images of HEK-293T cells expressing LIPA-α-Syn exposed to blue light for 24h (0.8 mW/mm²) and fixed with 4% PFA. Cells were immunolabeled with pS129 (Abcam, cyan) and stained with LipidSpot488 (yellow). (Scale bar = 2 µm). (**C**) Confocal images of induced aggregates in HEK-293T cells expressing LIPA-α-Syn exposed to blue light for 24h (0.8 mW/mm²) and fixed with 4% PFA. Cells were immunolabeled with pS129 (Abcam, cyan) and stained with LipidSpot488 (yellow) (Scale bar = 2 µm) and (**D**) the corresponding intensity profile of an irreversible aggregate engulfing a lipid droplet. (**E**) Confocal images of HEK-293T cells expressing LIPA-α-Syn (cyan) and stained for lipid droplets with LipidSpot488 (yellow), exposed to blue light with increasing durations (0, 3, 6, 12, and 24h). Nuclei are stained with DAPI (gray). (**F**) Quantification of the mean number of lipid droplets per cell (* = vs 0h) (N = 3). Data are presented as means ± SEM; \*\**p* ≤ 0.01; \*\*\**p* < 0.0001. (**G**) Quantification of the mean lipid droplet size (arbitrary units) (* = vs 0h) (N = 3). Data are presented as means ± SEM; \*\**p* ≤ 0.001. (**H**) Confocal images of HEK-293T cells expressing LIPA-α-Syn (magenta) and stained for lipid droplets LipidSpot488, (yellow) and pS129 (cyan). Cells were treated with DMSO or oleic acid (100 µM), exposed to blue light for 3h to induce reversible aggregates, and then fixed 24h post-illumination. Scale bar = 10 µm. (**I**) Quantification of the percentage of cells with aggregates relative to the total number of cells after cessation of illumination (* = vs DMSO) (N = 3). Data are presented as means ± SEM; ***p < 0.0001. (**J**) Confocal images of HEK-293T cells expressing LIPA-α-Syn (magenta) and stained for lipid droplets (LipidSpot, yellow) and pS129 (cyan). Cells were with DMSO or with both DGAT1 (10 µM) and DGAT2 (10 µM) inhibitors, exposed to blue light for 12h (irreversible aggregates), and fixed after 24h in the dark. (Scale bar = 10 µm). (**K**) Quantification of the percentage of cells with aggregates relative to the total number of cells after cessation of illumination (* = vs DMSO) (N = 3). Data are presented as means ± SEM; \*\*\*\**p* < 0.0001. (**L**) Representative confocal images of iNeurons expressing LIPA-α-Syn (magenta) and stained for lipid droplets (LipidSpot, yellow) and pS129 (cyan). Neurons were treated with DMSO or oleic acid (200 µM), exposed to blue light for 1.5h (reversible aggregates), then fixed immediately or after 6h and 24h post-illumination. (Scale bar = 5 µm). (**M**) Quantification of the percentage of neurons with aggregates relative to the total number of cells after cessation of illumination (* = vs DMSO) (N = 3). Data are presented as means ± SEM; \*\*\*\**p* < 0.0001.

Interestingly, analysis of the kinetics of annular LIPA-α-Syn structures formation during the transition from reversible to irreversible aggregation revealed a progressive increase in LDs accumulation. LDs were nearly undetectable before light stimulation but progressively increased throughout light exposure and LIPA-α-Syn aggregation (**Figure 4E**). Morphometric analysis across the aggregation time course demonstrated a marked significant, time-dependent increase in the number of LDs per cell, from approximately 0.5 at 0h to ∼2 at 6h, ∼5 at 12h, and ∼8 at 24h (**Figure 4E, F**). In parallel, the mean LD size also significantly increased progressively from early to late stages of aggregation, consistent with LD maturation and hypertrophy (**Figure 4G**). Together, these findings reveal a temporal association between α-Syn aggregation and progressive LD accumulation and enlargement, suggesting that α-Syn aggregation is accompanied by a sustained cellular lipid response. This lipid remodeling may contribute to the stabilization and persistence of α-Syn inclusions as aggregates transition toward an irreversible state.

To investigate whether LDs actively contribute to the stabilization of LIPA-α-Syn inclusions, we pharmacologically manipulated LD formation prior to light stimulation. Interestingly, treatment with oleic acid (OA), which promotes LD formation ^73^, significantly increased the persistence of otherwise reversible LIPA-α-Syn inclusions generated after 3h of light stimulation (**Figure 4H, I**). Conversely, inhibition of LD biogenesis using DGAT1/2 inhibitors converted the irreversible LIPA-α-Syn inclusions generated after prolonged light stimulation (12h) into a more reversible state, with inclusions rapidly and significantly dissipating within 24h following cessation of illumination (**Figure 4J, K**). Together, these findings support a model in which α-Syn aggregation is accompanied by LD accumulation and maturation, which in turn promotes the stabilization and persistence of α-Syn inclusions during their transition toward an irreversible state.

Notably, a similar effect was observed in iNeurons. OA treatment, which promotes LD accumulation, increased the persistence of early reversible LIPA-α-Syn aggregates, resulting in a significantly higher proportion of cells retaining inclusions following light cessation (**Figure 4L, M**). These findings indicate that the relationship between LD accumulation and α-Syn inclusion stabilization is conserved in human neuronal cells.

### Irreversible aggregates sensitize cells to stress and increase cell vulnerability

To confirm the pathophysiological relevance of our model, we assessed whether the transition to an irreversible, lipid-enriched state compromises cell viability. In neurodegenerative diseases, toxicity is often unmasked by a metabolic or environmental “second hit” (double-hit hypothesis) ^74^. We therefore exposed cells containing reversible or irreversible aggregates to a metabolic stress induced by nutrient deprivation (DMEM without FBS) for 4h, then analyzed survival and mitochondrial function ^40^. We first quantified cell survival by automated nuclear counting (DAPI). Wild type (WT) HEK-293T cells exhibited minimal loss of viability under nutrient deprivation, regardless of whether they were maintained in the dark or subjected to short (3h) or prolonged (24h) blue-light illumination (**Figure 5A**). In contrast, LIPA-α-Syn-expressing cells harboring irreversible aggregates displayed increased vulnerability even under nutrient-rich conditions, with cell survival significantly decreasing by approximately 30% over 24h (from 100% to ∼67%). This vulnerability was markedly exacerbated by nutrient deprivation, which resulted in approximately 70% cell loss (**Figure 5A**). To determine whether the observed toxicity was associated with impaired cellular metabolic activity, we performed an MTT assay (**Figure 5B**). In HEK-293T WT cells, MTT reduction remained relatively stable between 0 and 24h under both nutrient-rich and nutrient-deprived conditions, with no significant decrease in metabolic activity. In contrast, LIPA-α-Syn-expressing cells exhibited a marked and significant decline in metabolic activity even under control conditions, with an approximately 45% reduction over 24h. This impairment was further exacerbated by nutrient deprivation, resulting in a significant ∼60-65% reduction relative to early time points. Thus, cells harboring irreversible LIPA-α-Syn aggregates displayed a pronounced vulnerability to metabolic stress, whereas the same deprivation conditions were comparatively well tolerated by WT HEK-293T cells. The combination of irreversible aggregates and metabolic stress is therefore particularly deleterious, suggesting that persistent α-Syn aggregation compromises cellular metabolic capacity and reduces the ability of cells to adapt to conditions of increased metabolic demand. These results suggest that LDs-associated aggregates, compromise cellular metabolic activity and survival under stress conditions.

**Figure 5:**
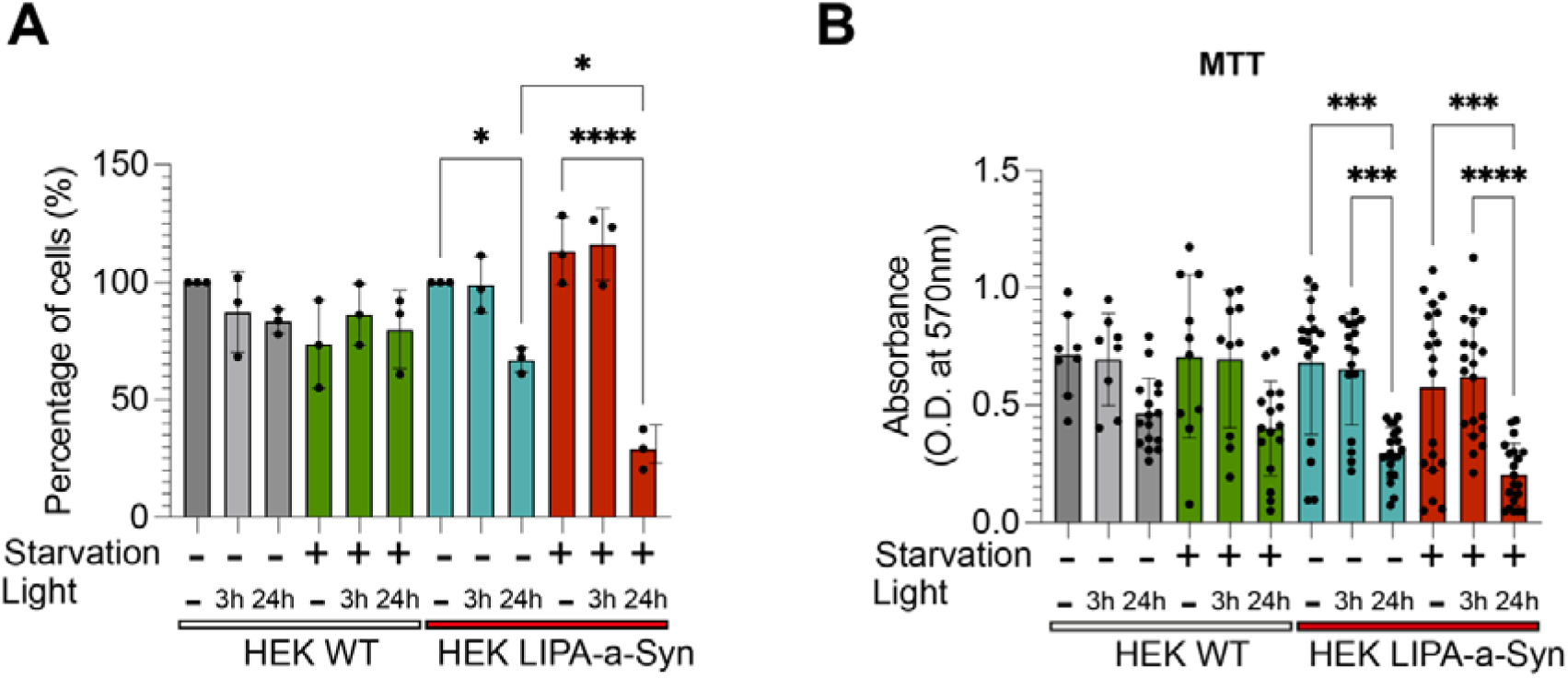
Irreversible aggregates sensitize cells to nutrient deprivation stress and induce mitochondrial dysfunction. (**A**) Quantification of cell number (DAPI nuclear counting) for wild-type (WT) HEK-293T cells or HEK-293T cells expressing LIPA-α-Syn, maintained in complete medium (−) or subjected to nutrient deprivation (DMEM without FBS, Starvation (+)) for 3h or 24h after aggregate induction. (* = vs control conditions) (N = 3). Data are presented as means ± SEM; \**p* ≤ 0.05; \*\*\*\**p* < 0.0001. (**B**) Quantification of mitochondrial metabolic activity by MTT assay (570 nm absorbance) in HEK-293T cells (WT or LIPA-α-Syn) in complete medium (−) or under nutrient deprivation (DMEM without FBS, (+)) at the indicated post-induction times (* = vs control conditions) (N = 3). Data are presented as means ± SEM; \*\*\**p* ≤ 0.001; \*\*\*\**p* < 0.0001.

## Discussion

For more than two decades, α-Syn aggregation has been tightly linked to dopaminergic neurodegeneration in PD, yet the fine spatiotemporal dynamics governing the emergence of LB-like inclusions remain incompletely understood ^75,76^. The central question we addressed here is: which mechanisms drive the transition from reversible condensates to structurally and biochemically irreversible inclusions? Conventional models, α-Syn PFFs or α-Syn overexpression have been instrumental for demonstrating seeding, spreading, and toxicity, but they offer limited temporal control over early aggregation steps. PFF paradigms bypass endogenous nucleation by introducing exogenous seeds. In contrast, viral overexpression yields slow, asynchronous aggregate formation, making it difficult to capture transient intermediates and maturation events ^48,49^. Using an optogenetic approach (LIPA-α-Syn), we sought to bridge this gap by following α-Syn as it evolves from a soluble monomer to pathological inclusions through a controlled, reversible state ^50^. This system provides a precise control over the onset and duration of aggregation, enabling systematic interrogation of maturation kinetics and supporting the emerging view that inclusions are not static end-points but dynamic entities that evolve in both physical state and molecular composition over time ^21^.

### A temporal phase transition between reversible and irreversible aggregates

A key conceptual advance of this work is the operational distinction between reversible and irreversible aggregates, not based on size or simple detergent insolubility, but on their biophysical properties, biochemical signatures, and fate under cellular quality control. Rather than a binary “soluble versus insoluble” model, our findings support a continuum of α-Syn states punctuated by a critical phase transition. By titrating the duration of light stimulation, we identified a reproducible temporal tipping point beyond which aggregates cease to be cleared and instead persist throughout the 24 h post-illumination period, in both HEK-293T cells and human neuronal culture. Below this threshold (short illumination), aggregates remain largely reversible, whereas longer exposures drive an abrupt shift to a stable, degradation-resistant state. Conceptually, this temporal threshold may correspond to a stage beyond which proteostasis systems, primarily the UPS, can no longer restore the native proteome ^62,63^. Similar ideas have been proposed for prion-like spreading, in which early inclusions may still be remodelled or removed, while late-stage assemblies become self-sustaining and propagation-competent ^77,78^.

### From liquid-like condensates to solid, Lewy body-like shells

Biophysically, our data refine the trajectory of α-Syn assemblies along the phase-separation landscape. Early aggregates formed after short illumination behave as dense, yet dynamic condensates: they retain high FRAP recovery, remain sensitive to 1,6-hexanediol, and are efficiently removed once the stimulus is withdrawn, features consistent with a liquid-like or gel-like LLPS state ^28,79^. These condensates display increased local concentration and partial resistance to mild detergent, but internal molecules remain mobile and predominantly held together by weak, multivalent interactions. Over time, however, we observe a progressive loss of molecular mobility, loss of hexanediol sensitivity, and emergence of a rigid, annular morphology by STED microscopy, with α-Syn forming shells around a central cavity. This is reminiscent of the “aging” of protein condensates described for other aggregation-prone proteins, in which liquid droplets gradually convert into gel-like or solid phases that could be enriched in β-sheet crosslinks ^22,29,80^.

This physical transition is tightly coupled to a stereotyped biochemical remodeling. We find that phosphorylation at S129, long recognized as a robust marker of Lewy pathology, becomes nearly ubiquitous at late time points, while tyrosine phosphorylations (Y39, Y125, Y133) accumulate more gradually ^30,31^. Importantly, pS129 in early reversible aggregates remains accessible to alkaline phosphatase, whereas in late solid inclusions it becomes resistant, consistent with steric occlusion within a compact structure ^65^. PLK inhibition with BI2536 markedly reduces pS129, while PLK2 overexpression increased aggregate stability, supporting a causal contribution of site-specific phosphorylation to the locking process ^33–35^. Paradoxically, the phospho-deficient mutant S129A also exhibited enhanced stability, although less pronounced compared to stability induced under PLK2-induced phosphorylation condition. This suggests that modifying the S129 residue by phosphorylation or mutation promotes aggregate stabilization. This effect is likely independent of normal protein behavior, as recently shown, the S129A mutation does not alter α-Syn expression levels or cell toxicity. These findings indicate that S129A is not a neutral loss-of-function mutant but instead may introduce structural changes that affect aggregate properties. In parallel, late aggregates progressively recruit ubiquitin and p62/SQSTM1, core components of LBs and other proteinopathies, echoing proteomic studies of human LB material ^14,81,82^. Thus, the LIPA system recapitulates both the physical aging and molecular “Lewy body signature” of α-Syn inclusions in a time-resolved manner.

### A Vicious Cycle Between Proteostasis Collapse and Irreversible α-Syn Aggregation

Our results underscore a bidirectional relationship between α-Syn maturation and cellular proteostasis. Inhibiting the UPS with MG132 or epoxomicin nearly abolishes clearance of otherwise reversible aggregates, whereas autophagy inhibition exerts only a partial effect, indicating that the proteasome constitutes the first line of defense against early condensates ^83,84^. Conversely, once aggregates reach the solid, Lewy-like state, two independent UPS reporters (YFP-CL1 and Ub-R-YFP) accumulate robustly, to levels comparable with or even exceeding, pharmacological proteasome blockade ^58^. These observations are consistent with prior evidence that misfolded or fibrillar α-Syn can impair UPS activity and foster accumulation of ubiquitinated proteins ^85–88^.

Together, these data support a vicious-cycle model: modest UPS dysfunction slows aggregate clearance and favors their maturation into rigid inclusions; in turn, these inclusions further saturate and inhibit UPS capacity, driving a self-reinforcing proteostatic catastrophe ^89,90^. Importantly, in our system this cycle is largely reversible at early time points: when aggregates are still liquid-like, proteasome function is sufficient to restore the proteome once the aggregation trigger is removed. Only after crossing the maturation threshold does the system become trapped in a high-burden state in which both inclusions and proteostasis defects mutually amplify.

### Lipid droplets as modulators of α-Syn condensate maturation

An emerging concept supported by our findings is that LDs are not inert bystanders but active structural modulators that shape α-Syn condensate maturation. Using LipidSpot staining and super-resolution imaging, we reveal that late, irreversible aggregates consistently adopt an annular donut-like architecture in which phosphorylated α-Syn tightly surrounds a neutral lipid droplet. This organization parallels recent ultrastructural analyses of human LBs and experimental models, which describe inclusions rich in vesicles, fragmented organelles, and lipid material rather than homogeneous fibrillar cores ^14,37,81^.

Functionally, LDs and aggregates engage in a reciprocal relationship. On the one hand, α-Syn aggregation triggers robust LD increased abundance and hypertrophy over time, consistent with broader evidence that proteotoxic or metabolic stress can drive LD formation via mTORC1-SREBP signaling and DGAT-dependent triglyceride synthesis ^38–40,91^. On the other hand, experimentally increasing LD abundance with oleic acid is sufficient to convert normally reversible condensates into irreversible inclusions in both HEK-293T cells and human neurons, whereas inhibiting DGAT1/2 partially restores reversibility even after prolonged induction. These results align with recent work showing that LDs promote aberrant LLPS and aggregation of α-Syn, acting as scaffolds that concentrate the protein and alter its conformational landscape ^38,44,92^. Consistent with our observations, Cevallos et al., recently showed that increasing LD abundance promoted the formation of LD-rich α-Syn condensate, supporting a role for LDs as drivers of aberrant phase separation ^44^. The amphipathic N-terminus of α-Syn has a high affinity for curved lipid interfaces, and heterogeneous nucleation at LD surfaces likely accelerates the transition from liquid-like droplets to solid shells ^37,44,92^.

This architecture may have important metabolic consequences. By sequestering LDs within α-Syn annular assemblies, irreversible aggregates could perturb the organization of LD-associated proteins and impair LD turnover, which could limit the mobilization of LD-derived fatty acids that support mitochondria energy production during cellular stress ^39,91,93^. In our nutrient-deprivation paradigm, cells harboring irreversible, LD-encapsulating aggregates exhibit profound mitochondrial failure and massive cell loss, whereas aggregate-free controls tolerate the same challenge with minimal impact. Given the extraordinary energetic demands of dopaminergic neurons, which rely on flexible fuel usage and intact mitochondrial networks, such LD sequestration could be a key driver of their selective vulnerability ^94,95^.

### An integrated sequential model of Lewy body biogenesis

Integrating these findings, we propose a sequential and mechanistically explicit model of LB biogenesis. First, α-Syn undergoes LLPS to form dynamic condensates that, while locally concentrated and partially detergent-resistant, remain liquid- or gel-like, proteasome-accessible, and largely reversible upon stimulus withdrawal. Second, with time and in the presence of sufficient lipid droplet scaffolds and kinase activity, these condensates maturate into annular shells that encapsulate LDs, acquire a dense constellation of pathological PTMs (pS129, pY39, pY125, pY133), and recruit ubiquitin/p62. This transition coincides with steric protection of pS129 from dephosphorylation and a shift from weak, reversible interactions toward more stable and less dynamic assemblies. Third, once this structural and biochemical threshold is crossed, inclusions become functionally irreversible: they resist cellular clearance, saturate the UPS, sequester metabolic resources, and dramatically sensitize cells to otherwise sublethal metabolic stress, culminating in mitochondrial collapse and cell death.

### Therapeutic implications: targeting the phase of α-Syn

Our work has several therapeutic implications. By defining a temporal window during which aggregates are still liquid-like or gelled and remain under proteostatic control, it suggests that interventions acting early on maturation, rather than solely on fibrillar end-products, may be most effective. In principle, three classes of strategies could be particularly attractive: (i) modulation of kinase pathways (e.g., PLK2, c-Abl) to limit pathological phosphorylation and structural locking ^33,96,97^; (ii) pharmacological manipulation of lipid metabolism, such as DGAT inhibition, SREBP modulation, or selected lipid-lowering agents, to reduce pathological LD scaffolding; and (iii) enhancement of proteostasis via UPS support, chaperone induction, or autophagy activation to maintain condensates within a reversible regime ^89,91,98,99^. By analogy with tafamidis stabilizing transthyretin tetramers in amyloidosis, one might envision a “pharmacology of phase” aimed at keeping α-Syn in a condensed but reversible state, forestalling its progression to lipid-encapsulated, Lewy body-like inclusions and thereby slowing PD progression ^100,101^.

## Supplemental material

**Supplemental Figure S1:**
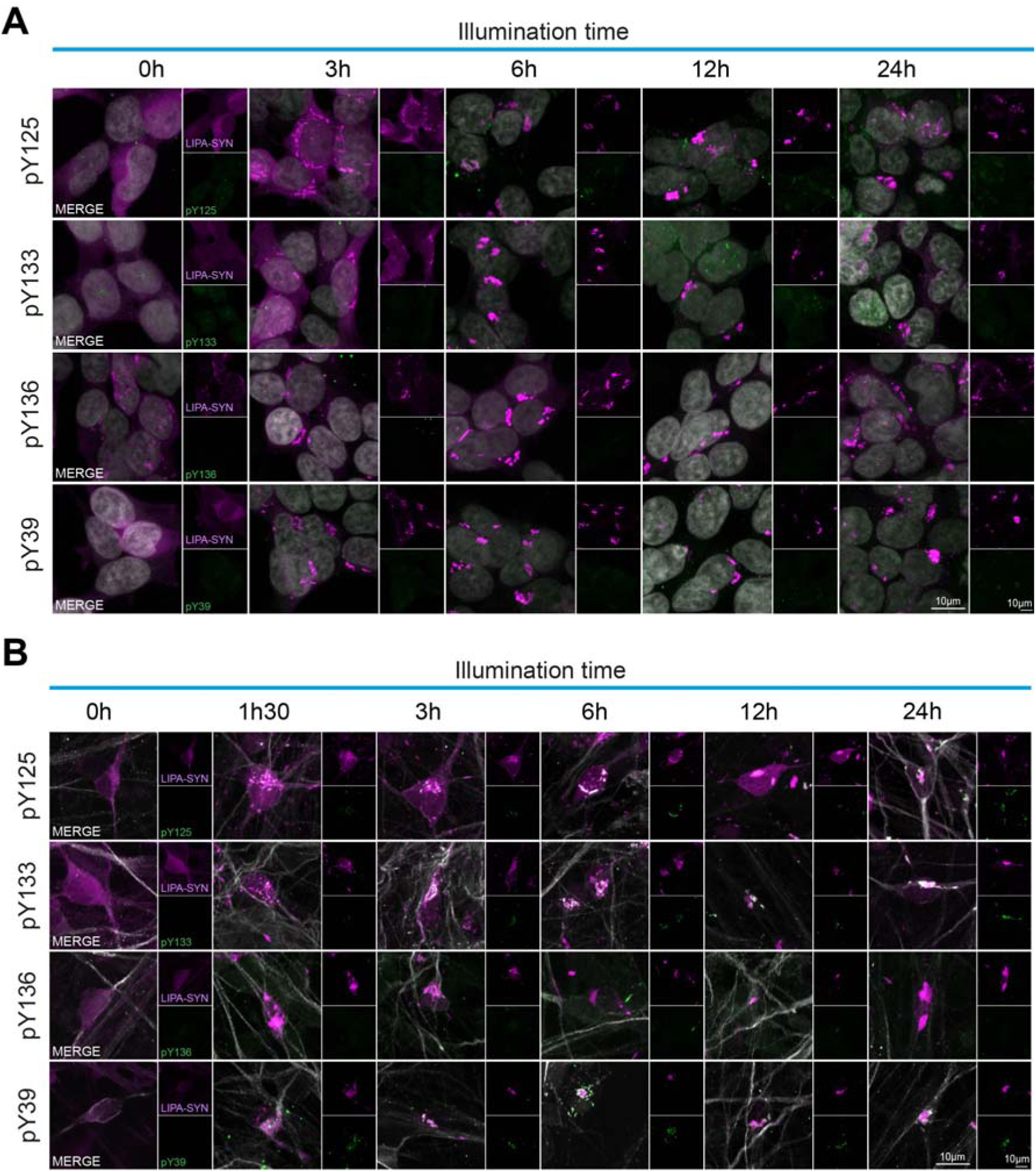
The transition of LIPA-α-Syn aggregates from a reversible to an irreversible state is associated with specific PTMs. (**a**) Confocal images of HEK-293T cells expressing LIPA-α-Syn (magenta) exposed to blue light for 3, 6, 12, or 24h (0.8 mW/mm²) or not exposed (0h) and fixed with 4% PFA. Cells were immunolabeled for different PTMs (green): pY125, pY133, pY136, and pY39. (Scale bar = 10 µm). (**b**) Confocal images of iNeurons expressing LIPA-α-Syn (magenta) exposed to blue light for 1.5, 3, 6, 12, or 24h (0.1 mW/mm²) or not exposed (0h) and fixed with 4% PFA. Neurons were immunolabeled for different PTMs (green): pY125, pY133, pY136, and pY39. (Scale bar = 10 µm).

**Supplemental Figure S2:**
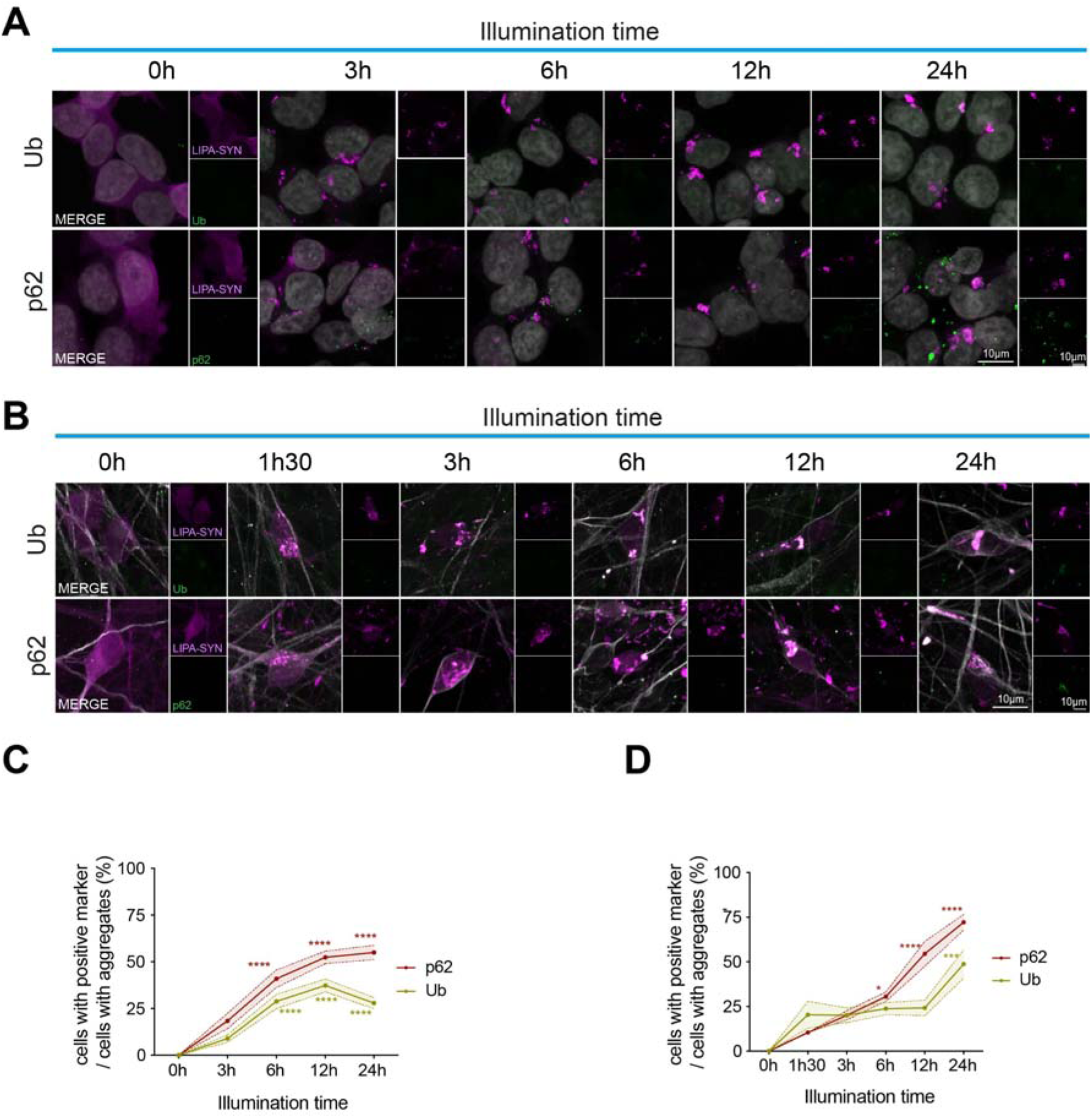
The transition of LIPA-α-Syn aggregates from a reversible to an irreversible state is associated with the accumulation of specific Lewy Body-associated markers. (**A**) Confocal images of HEK-293T cells expressing LIPA-α-Syn (magenta) exposed to blue light for 3, 6, 12, or 24h (0.8 mW/mm²) or not exposed (0h) and fixed with 4% PFA. Cells were immunolabeled for different LB markers (green): ubiquitin (Ub) and p62/SQSTM1 (p62). (Scale bar = 10 µm). (**B**) Confocal images of iNeurons expressing LIPA-α-Syn (magenta) exposed to blue light for 1.5, 3, 6, 12, or 24h (0.1 mW/mm²) or not exposed (0h) and fixed with 4% PFA. Neurons were immunolabeled for different LB markers (green): ubiquitin (Ub) and p62/SQSTM1 (p62). (Scale bar = 10 µm). (**C**) Quantification of the percentage of LIPA-α-Syn aggregate-positive cells for each Lewy body-associated marker relative to the total number of cells containing aggregates. Values were compared with those for the 3h condition, which represents the most reversible aggregates. (* = vs 3h) (N = 3). Data are presented as means ± SEM; *p ≤ 0.05; **p ≤ 0.01; ***p ≤ 0.001; ****p < 0.0001. (**D**) Quantification of the percentage of LIPA-α-Syn aggregate-positive iNeurons for each Lewy body-associated marker relative to the total number of cells containing aggregates. Values were compared with the 1.5h condition, which represents the most reversible aggregates. (* = vs 1.5h) (N = 3). Data are presented as means ± SEM; *p ≤ 0.05; **p ≤ 0.01; ***p ≤ 0.001; ****p < 0.0001.

**Supplemental Figure S3:**
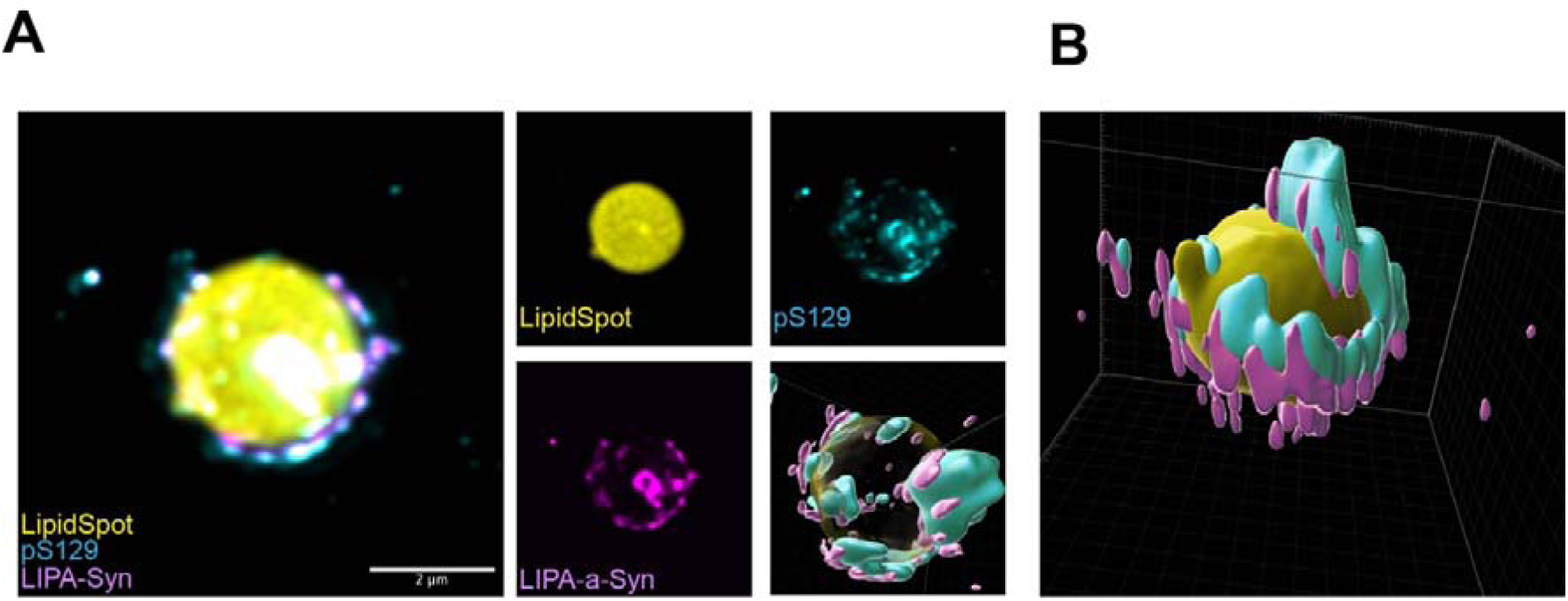
Irreversible aggregates display a hollow-centered morphology that engulfs lipid droplets. (**A**) STED images of induced aggregates in HEK-293T cells expressing LIPA-α-Syn exposed to blue light for 24 h (0.8 mW/mm²) and fixed with 4% PFA. Cells were immunolabeled with pS129 (Abcam, cyan) and stained with LipidSpot (yellow). (Scale bar = 2 µm). (**B**) Imaris 3D rendering of an irreversible aggregate engulfing a lipid droplet.

